# Soluble TREM2 reduces DAP12 surface expression by dissociating the TREM2–DAP12 complex

**DOI:** 10.64898/2026.05.05.723083

**Authors:** Atsuma Yamada, Shori Yatame, Fuminori Tsuruta

## Abstract

Triggering receptor expressed on myeloid cells 2 (TREM2) plays a crucial role in regulating various microglial functions, including phagocytosis, inflammation, chemotaxis, and proliferation. Recent studies have demonstrated that TREM2 associates with DAP12 to mediate intracellular signaling essential for these processes. Despite the importance of the TREM2–DAP12 complex in microglial physiology, the mechanisms controlling its expression and activity remain poorly understood. In this study, we show that the soluble ectodomain of TREM2 (sTREM2) regulates microglial phagocytic activity by attenuating the surface expression of DAP12. Stimulation with recombinant sTREM2 reduced the surface expression of DAP12, but not that of TREM2. In addition, sTREM2 bound to full-length TREM2, leading to the uncoupling of TREM2 from DAP12. Furthermore, pretreatment of microglial cell line BV2 with sTREM2 significantly inhibited amyloid-β uptake. These findings suggest that sTREM2 negatively regulates TREM2 signaling by destabilizing the TREM2–DAP12 complex and may act as a novel modulator of TREM2 signaling under physiological and pathological conditions.

## Introduction

Microglia are the resident immune cells of the central nervous system (CNS) and play fundamental roles in maintaining brain homeostasis (1). Microglia arise from yolk sac progenitors during early embryogenesis, colonize the brain prior to vascular maturation, and persist throughout life via self-renewal (2, 3). In the healthy brain, microglia dynamically survey their environment, sensing neuronal activity and extracellular signals (4–6). Microglia also regulate neurodevelopment by shaping synapse formation and elimination (7–10). Thus, microglia are critical for normal brain development. In contrast, microglia function as key immune mediators in disease. In response to injury, stress, infection, or environmental changes, microglia adopt reactive states accompanied by alterations in morphology, gene expression, and function, including the release of inflammatory mediators (11). Although these responses are initially neuroprotective, their dysregulation can drive disease progression (12, 13). Emerging evidence further indicates that microglial properties are highly context-dependent and change across the lifespan, with aging-associated shifts frequently linked to impaired homeostasis and a heightened inflammatory bias (14). Such alterations in microglial function are increasingly recognized as key contributors to neurological and psychiatric disorders, highlighting their importance in both disease progression and brain aging.

Triggering receptor expressed on myeloid cells 2 (TREM2) is a microglia-enriched cell-surface receptor that plays a central role in innate immune sensing in the central nervous system (15). TREM2 signaling is mediated through its association with the adapter protein DNAX-activation protein 12 (DAP12) encoded by the *Tyrobp* gene. Upon ligand binding to TREM2, the immunoreceptor tyrosine-based activation motif within the cytoplasmic domain of DAP12 is phosphorylated by Src family kinases. This phosphorylation recruits the protein tyrosine kinase, SYK, which activates downstream signaling cascades such as the phosphoinositide 3-kinase pathway (16, 17). TREM2 recognizes a wide variety of ligands, including lipids, apolipoproteins, and cellular debris, thereby regulating microglial survival, metabolism, and phagocytic activity (17–22). Under physiological conditions, TREM2 contributes to tissue homeostasis by promoting the clearance of apoptotic cells (22, 23) and protein aggregates while limiting excessive inflammatory responses. In pathological conditions, TREM2 is required for the transition of microglia into disease-associated states that target specific sites like amyloid plaques (24, 25). Importantly, rare variants of TREM2 are linked to a higher risk of brain diseases, such as Alzheimer’s disease (AD) (26). Moreover, as microglia exhibit reduced metabolic capacity and diminished responsiveness to environmental stimuli with aging, TREM2 is thought to potentially contribute to these phenomena (27, 28). Ultimately, impaired TREM2 signaling leads to defective debris clearance and heightened neuronal vulnerability, proving that TREM2 is a critical regulator of microglial adaptation throughout the lifespan.

Soluble TREM2 (sTREM2) is generated by proteolytic shedding of the full-length receptor, mediated by the metalloproteinases a disintegrin and metalloproteinase 10 (ADAM10) and a disintegrin and metalloproteinase 17 (ADAM17). This cleavage occurs at a membrane-proximal site within the extracellular domain, between His157 and Ser158, and releases the ectodomain into the extracellular space (29–31). Emerging evidence suggests that sTREM2 is not merely a byproduct of receptor turnover but a biologically active molecule that modulates microglial homeostasis. Indeed, recent studies have shown that sTREM2 regulates key cellular processes, including microglial survival and inflammatory responses (32). In addition, soluble receptor fragments can influence the stability and signaling of their membrane-bound counterparts, raising the possibility that sTREM2 modulates TREM2–DAP12 function (33). However, the precise molecular mechanisms underlying these effects—particularly whether sTREM2 directly interacts with cell-surface TREM2—remain largely unclear and warrant further investigation.

In this study, we identify a previously unrecognized mechanism by which extracellular sTREM2 regulates the assembly and membrane localization of the TREM2–DAP12 complex in the microglial cell line BV2. Treatment with sTREM2 selectively reduces the membrane localization of DAP12 relative to its total expression without altering TREM2 membrane localization. Moreover, sTREM2 interacts with the extracellular domain of TREM2, but not with DAP12, and weakens the TREM2–DAP12 interaction. Consistent with these observations, sTREM2 treatment suppresses the phagocytic uptake of substrates, including amyloid-β. Collectively, these findings support a model in which sTREM2 binds to TREM2 and destabilizes the TREM2–DAP12 complex, thereby modulating microglial function.

## Results

### sTREM2 decreases surface expression of DAP12

TREM2 is synthesized in the endoplasmic reticulum and is subsequently transported to the plasma membrane. Upon forming a complex with DAP12, TREM2 is partially cleaved by ADAM10 and ADAM17, releasing sTREM2 into the extracellular space (Fig. 1A). Importantly, disease-associated TREM2 variants exhibit impaired trafficking to the plasma membrane, leading to reduced surface expression and sTREM2 production (34). This observation suggests that TREM2-mediated pathological changes are tightly regulated by the subcellular localization of TREM2. Therefore, we first investigated whether sTREM2 regulates the surface expression levels of TREM2. To this end, we purified His-tagged recombinant sTREM2 protein (amino acids 1–157) from the conditioned medium of transfected HEK293T cells and performed a cell-surface biotinylation assay (Fig. 1B to 1D). Our analysis revealed that pretreatment with sTREM2 did not significantly alter the levels of biotinylated TREM2 on the plasma membrane (Fig. 1E and 1F). Intriguingly, however, we observed a significant reduction in the surface expression of DAP12, a key signaling partner of TREM2 (Fig. 1G and 1H). These findings suggest that sTREM2 stimulation selectively triggers the internalization or depletion of DAP12, rather than affecting TREM2 localization itself. To further confirm this effect, we performed immunocytochemistry for either TREM2 or DAP12 in the presence or absence of cell permeabilization (29, 35) (Fig. 1I). Consistent with the biotinylation assay results, surface TREM2 levels remained unchanged (Fig. 1J and 1K). However, pretreatment with sTREM2 significantly decreased the cell-surface expression of DAP12 (Fig. 1L and 1M), indicating that sTREM2 selectively reduces DAP12 levels on the plasma membrane. To determine whether the reduction in cell-surface DAP12 results from internalization or impaired trafficking to the plasma membrane, we performed the same immunocytochemistry at 4°C, at which endocytic processes are suppressed. Although sTREM2 reduced the cell-surface expression of DAP12 at 37°C, this effect was attenuated at 4°C (Fig. 1N and 1O). These findings raise the possibility that the reduction in cell-surface DAP12 involves an internalization process.

**Figure 1.**
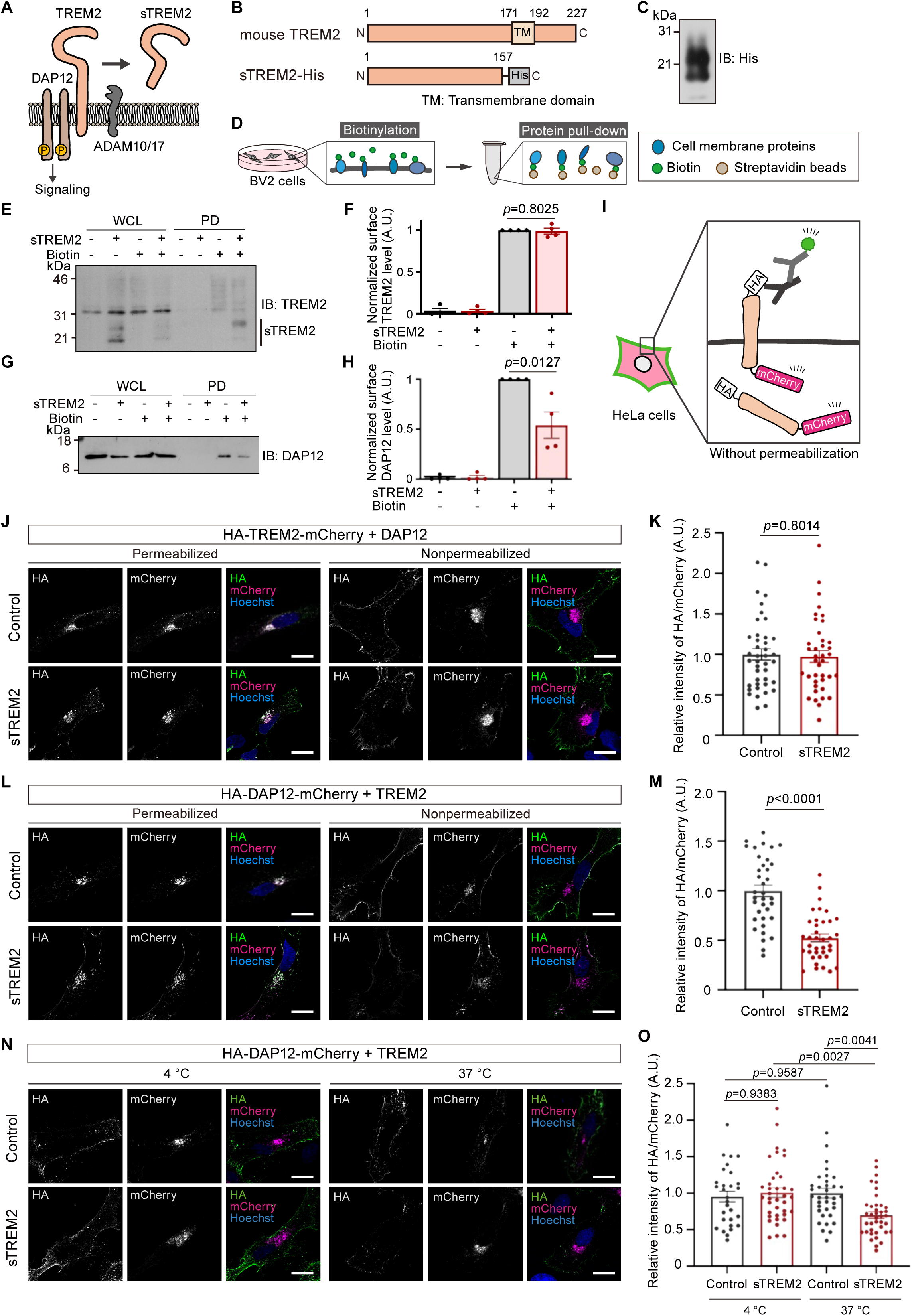
sTREM2 decreases surface expression of DAP12. **(A)** Schematic illustration of the formation and shedding of the TREM2–DAP12 complex at the cell surface. TREM2 interacts with DAP12 via their respective transmembrane domains, where phosphorylation of DAP12 triggers downstream signaling. TREM2 undergoes proteolytic cleavage, primarily mediated by ADAM10, to release soluble TREM2 (sTREM2). **(B)** Schematic structure of TREM2 and sTREM2. **(C)** Immunoblotting of recombinant sTREM2-His protein purified from HEK293T conditioned medium. **(D)** Schematic illustration of the cell-surface biotinylation assay. Cell surface proteins were labeled with biotin and pulled down using streptavidin beads. **(E)** Immunoblot analysis of biotinylated cell surface TREM2. **(F)** Relative quantification of surface TREM2. The bar graph shows the ratio of precipitated TREM2 (PD) to TREM2 levels in whole cell lysates (WCL). sTREM2 treatment did not affect the total expression levels of TREM2. PD, pull-down; WCL, whole cell lysate. n=4. Data are presented as mean ± SEM. Student’s *t*-test. **(G)** Immunoblot analysis of biotinylated cell surface DAP12. **(H)** Relative quantification of surface DAP12. The bar graph shows the ratio of precipitated DAP12 (PD) to DAP12 levels in whole cell lysates (WCL). sTREM2 decreased surface expression of DAP12. n=4. Data are presented as mean ± SEM. Student’s *t*-test. **(I)** Schematic illustration of the cell surface localization assay. The N-terminal HA tag is detected exclusively at the cell surface under non-permeabilized conditions, whereas the C-terminal mCherry signal represents total protein expression. **(J)** Representative images of HA-TREM2-mCherry expressed in HeLa cells. Scale bar, 20 µm. **(K)** HA signal intensities at the plasma membrane were quantified and normalized to mCherry fluorescence to assess relative surface expression levels. Control, n=41; sTREM2, n=39. Data are presented as mean ± SEM. Student’s *t*-test. **(L)** Representative images of HA-DAP12-mCherry expressed in HeLa cells. Scale bar: 20 µm. **(M)** HA signal intensities at the plasma membrane were quantified and normalized to mCherry fluorescence to assess relative surface expression levels. Control, n=36; sTREM2, n=37. Data are presented as mean ± SEM. Student’s *t*-test. **(N)** Representative images of HA-DAP12-mCherry expressed in HeLa cells. Scale bar, 20 µm. **(O)** HA signal intensities at the plasma membrane were quantified and normalized to mCherry fluorescence to assess relative surface expression levels. 4 °C: Control, n = 29; sTREM2, n = 38. 37°C: Control, n = 36; sTREM2, n = 39. Data are presented as mean ± SEM. Student’s *t*-test.

### sTREM2 interacts with TREM2

To examine whether sTREM2 interacts with DAP12, we performed a co-immunoprecipitation assay. We co-transfected HEK293T cells with expression plasmids for sTREM2-HA, TREM2-Flag, and DAP12-Flag, followed by immunoprecipitation with an anti-Flag antibody. While sTREM2-HA co-precipitated with TREM2-Flag, no interaction was detected with DAP12-Flag (Fig. 2A). This physical association contrasts with our functional observations, in which sTREM2 reduced the surface expression of DAP12, but had no effect on TREM2 levels. To further examine these findings, we performed a reciprocal immunoprecipitation using an anti-HA antibody and found that TREM2-His was co-precipitated with sTREM2-HA (Fig. 2B). These results further support the physical interaction between sTREM2 and TREM2. Collectively, our data suggest that sTREM2 preferentially associates with TREM2 rather than DAP12. Next, to identify the binding site, we generated two TREM2 mutants: one consisting solely of the extracellular Ig-like (Igl) domain (aa 1–130) and another lacking the Igl domain (amino acid Δ26–122; ΔIgl) (Fig. 2C). Our binding assays revealed that sTREM2 specifically interacts with the Igl domain (Fig. 2D). To further validate this, we performed co-immunoprecipitation using alternative epitope tags, which confirmed that Igl domains associate with each other (Fig. 2E). These results suggest that sTREM2 associates with TREM2 via the Igl domain.

**Figure 2.**
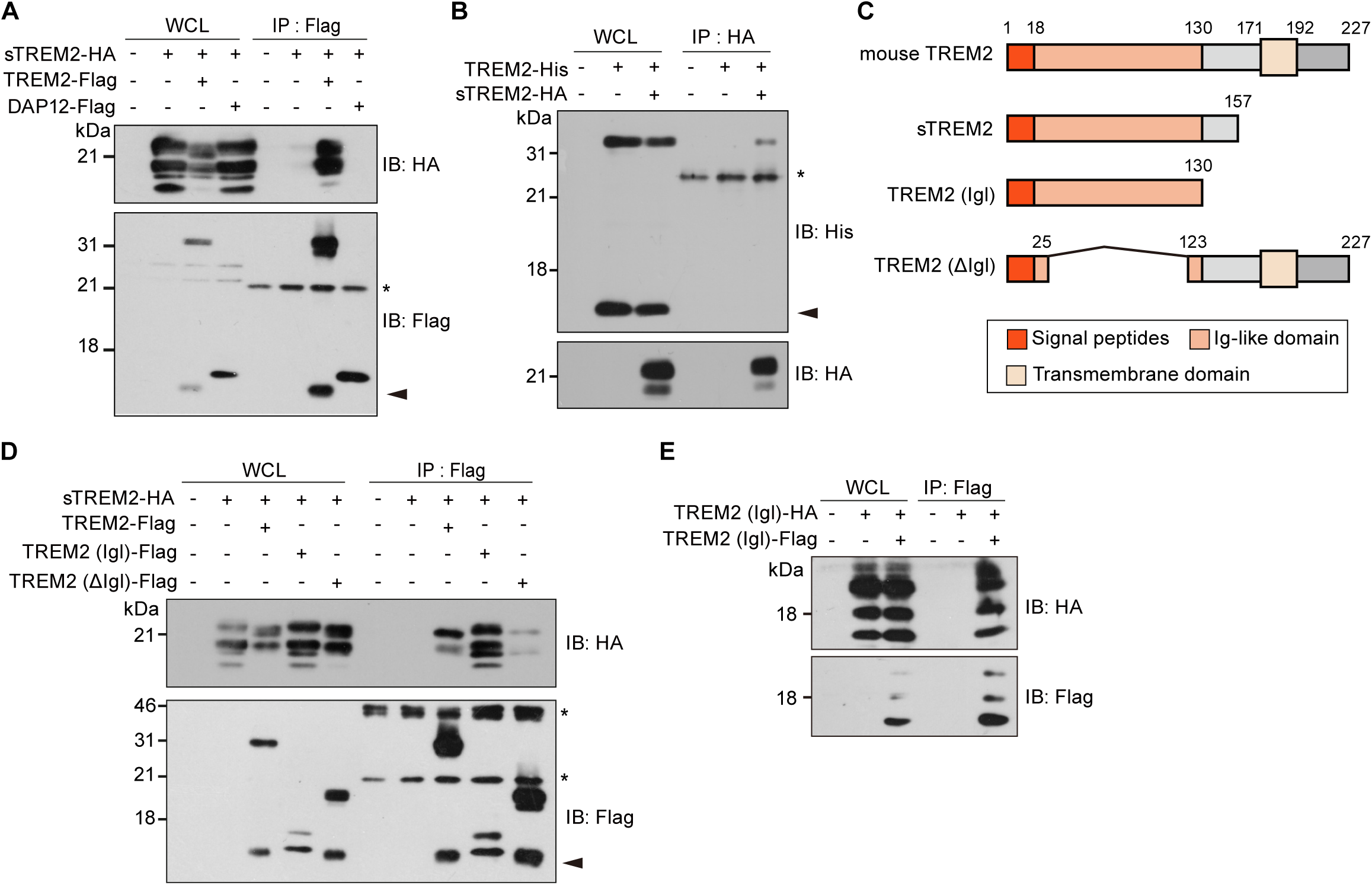
sTREM2 interacts with TREM2. **(A)** Co-immunoprecipitation of sTREM2-HA with either TREM2-Flag or DAP12-Flag in HEK293T cells. Co-precipitated proteins were analyzed using anti-HA and anti-Flag antibodies. Asterisk, nonspecific bands. Arrowhead, C-terminal fragments generated by proteolytic processing. **(B)** Co-immunoprecipitation of sTREM2-HA with TREM2-His in HEK293T cells. Co-precipitated proteins were analyzed by immunoblotting using anti-His and anti-HA antibodies. Asterisk, nonspecific bands. Arrowhead, C-terminal fragments generated by proteolytic processing. **(C)** Schematic structure of TREM2, sTREM2, the TREM2 Ig-like domain, and a TREM2 deletion mutant. **(D)** Co-immunoprecipitation of sTREM2-HA with Flag-TREM2 variants in HEK293T cells. Co-precipitated proteins were analyzed by immunoblotting using anti-Flag and anti-HA antibodies. Asterisk, nonspecific bands. Arrowhead, C-terminal fragments generated by proteolytic processing. **(E)** Co-immunoprecipitation of HA- and Flag-tagged TREM2 Igl constructs in HEK293T cells. Co-precipitated proteins were analyzed by immunoblotting with anti-HA and anti-Flag antibodies.

### sTREM2 impairs TREM2–DAP12 complex formation

Given that sTREM2 reduced the surface expression of DAP12 through its interaction with TREM2, we next investigated whether sTREM2 affects the formation of the TREM2–DAP12 complex. The association between TREM2 and DAP12 was markedly reduced upon co-expression of sTREM2 (Fig. 3A), indicating that sTREM2 interferes with the formation of the TREM2–DAP12 complex. To further confirm whether the sTREM2-induced dissociation of the TREM2–DAP12 complex underlies the reduction in DAP12 surface localization, we employed a TREM2 K183A mutant (Fig. 3B). K183 in mouse TREM2, corresponding to K186 in human TREM2 (hTREM2), is a transmembrane residue critical for the interaction with DAP12, and mutation of the corresponding residue to alanine disrupts TREM2–DAP12 complex formation (36). We first verified the binding deficiency of the K183A mutant via co-immunoprecipitation; while TREM2 WT robustly associated with DAP12, the K183A mutant failed to bind DAP12 (Fig. 3C). Using this construct, we assessed the surface expression of DAP12 using a cell-surface biotinylation assay. In cells expressing the K183A mutant, DAP12 surface expression was significantly lower than in cells expressing WT TREM2 (Fig. 3D and 3E). In addition, immunocytochemical analysis revealed that sTREM2 treatment did not significantly affect the surface expression of DAP12 in cells expressing the K183A mutant (Fig. 3F and 3G). Taken together, these findings suggest that sTREM2 promotes the dissociation of the TREM2–DAP12 complex through its interaction with TREM2, thereby reducing DAP12 surface expression.

**Figure 3.**
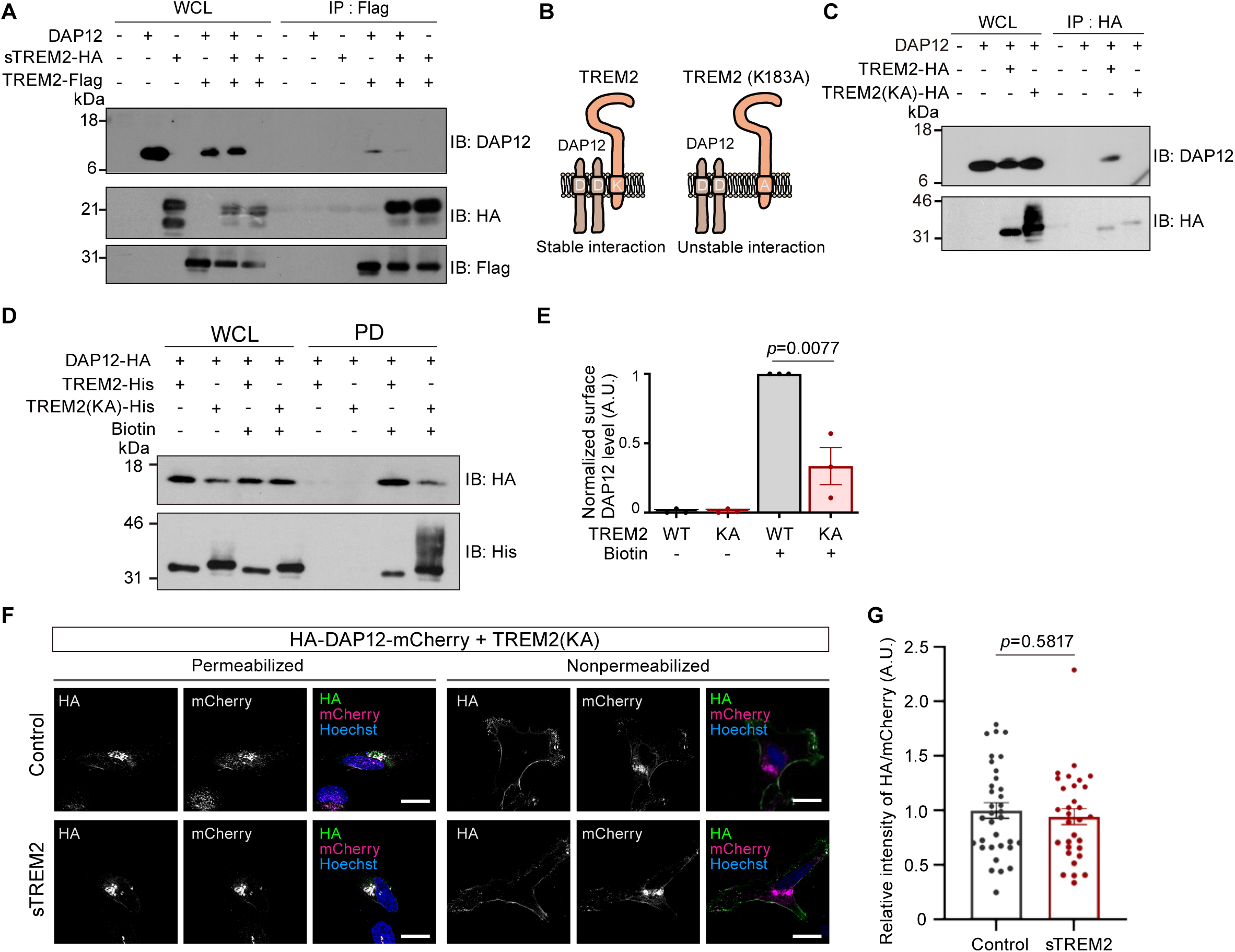
sTREM2 impairs TREM2–DAP12 complex formation. **(A)** Co-immunoprecipitation of DAP12, sTREM2-HA and TREM2-Flag in HEK293T cells. Co-precipitated proteins were analyzed using anti-DAP12, anti-HA and anti-Flag antibodies. **(B)** Schematic representation of the interaction between the transmembrane domains of TREM2 and DAP12. A lysine residue in TREM2 and an aspartic acid residue in DAP12 are critical for this interaction. Substitution of the lysine residue in TREM2 with alanine [TREM2 K183A (KA)] markedly reduces the interaction. **(C)** Co-immunoprecipitation of DAP12 with TREM2-HA or TREM2 (KA)-HA in HEK293T cells. Co-precipitated proteins were analyzed by immunoblotting with anti-DAP12 and anti-HA antibodies. **(D)** Immunoblot analysis of biotin-labeled cell-surface DAP12 in HeLa cells. **(E)** Relative quantification of surface DAP12. The bar graph shows the ratio of precipitated DAP12 (PD) to DAP12 levels in whole cell lysates (WCL). n=3. Data are presented as mean ± SEM. Student’s *t*-test. **(F)** Representative images of HA-DAP12-mCherry expressed in HeLa cells. Scale bar: 20 µm. **(G)** HA signal intensities at the plasma membrane were quantified and normalized to mCherry fluorescence to assess relative surface expression levels. Control, n=34; sTREM2, n=31. Data are presented as mean ± SEM. Student’s *t*-test.

### The effects of sTREM2 on TREM2 and DAP12 are conserved in humans

To determine whether the findings obtained in the mouse system are conserved in humans, we first compared the Igl domains of mouse and human sTREM2 (hsTREM2). The two domains shared 75.9% amino acid identity (Fig. 4A). Consistent with this sequence conservation, the AlphaFold 3-predicted structures exhibited highly similar overall folds, with a Cα root-mean-square deviation (RMSD) of 0.47 Å over 112 Cα atoms and a TM-score of 0.99 (Fig. 4B). Given the high conservation of the Igl domain, we next examined whether hsTREM2 interacts with hTREM2 or human DAP12 (hDAP12) by co-immunoprecipitation. We co-transfected HEK293T cells with expression plasmids for hsTREM2-HA, hTREM2-Flag, and hDAP12-Flag, followed by immunoprecipitation with an anti-Flag antibody. Whereas hsTREM2-HA co-precipitated with hTREM2-Flag, no interaction was detected with hDAP12-Flag (Fig. 4C). Reciprocal immunoprecipitation using an anti-HA antibody further confirmed the interaction between hsTREM2 and hTREM2 (Fig. 4D). Lastly, to examine the functional effect of hsTREM2, we purified recombinant hsTREM2 from the conditioned medium of HEK293T cells (Fig. 4E). Immunocytochemical analysis revealed that treatment with hsTREM2 significantly reduced the surface expression of hDAP12 (Fig. 4F and 4G). Together, these results indicate that the interaction between sTREM2 and TREM2 and the sTREM2-mediated reduction in DAP12 surface expression are conserved between mouse and human.

**Figure 4.**
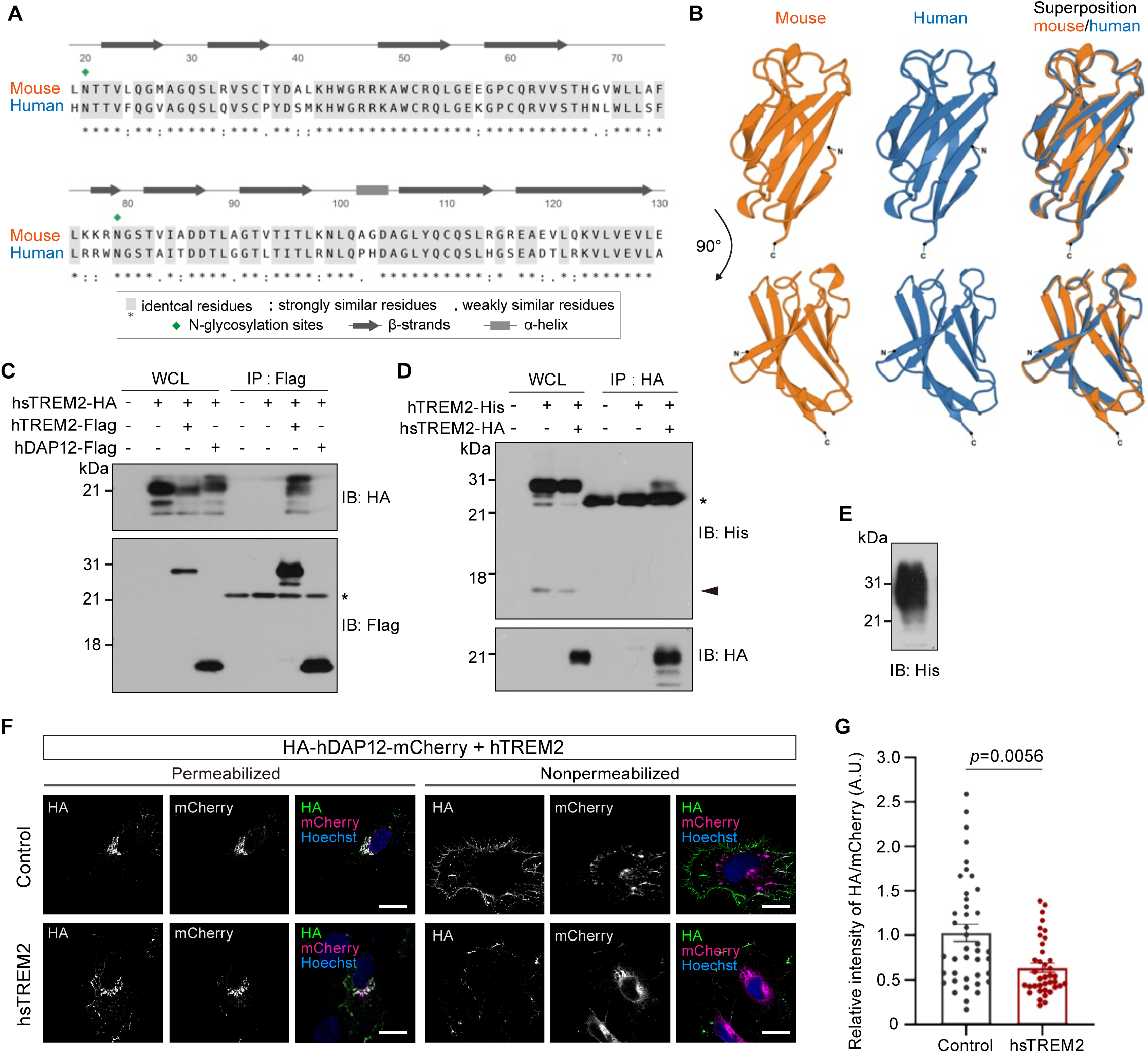
The effects of sTREM2 on TREM2 and DAP12 are conserved in humans. (A) The sequence alignment of mouse Igl domain and human Igl domain of sTREM2. The two sequences align without gaps and show 75.9% sequence identity (85/112 residues). Gray shading and asterisks, identical residues; colons, strongly similar residues. periods, weakly similar residues. diamonds, N-glycosylation sites; arrows, β-strands; box, α-helix. (B) AlphaFold 3 models of the Igl domain (aa 19–130) of mouse (orange) and human (blue) sTREM2 shown side by side in the same orientation (top) and rotated by 90° about the vertical axis (bottom). Mean pLDDT of the Igl domain (aa 19–130) was 92.3 for mouse and 96.2 for human. Structural superposition is shown on the right (cartoon representation). The Igl domains superimpose with a Cα RMSD of 0.47 Å over 112 Cα atoms and a TM-score of 0.99. Across all 25 pairwise comparisons between human and mouse models, the RMSD was 0.47 ± 0.09 Å (mean ± SD; range, 0.34–0.68 Å), comparable to the variation among models of the same species (human, 0.32 ± 0.18 Å; mouse, 0.18 ± 0.04 Å). The C-terminal stalk (aa 131–157) showed low prediction confidence (mean pLDDT, 48.3 for mouse and 51.6 for human), predicted to be disordered and was therefore omitted from the structural comparison. N and C indicate the N-and C-termini of the Igl domain. **(C)** Co-immunoprecipitation of human sTREM2 (hsTREM2)-HA with either human TREM2 (hTREM2)-Flag or human DAP12 (hDAP12)-Flag in HEK293T cells. Co-precipitated proteins were analyzed using anti-HA and anti-Flag antibodies. Asterisk, nonspecific bands. **(D)** Co-immunoprecipitation of hsTREM2-HA with hTREM2-His in HEK293T cells. Co-precipitated proteins were analyzed by immunoblotting using anti-His and anti-HA antibodies. Asterisk, nonspecific bands. Arrowhead, C-terminal fragments generated by proteolytic processing. **(E)** Immunoblotting of recombinant hsTREM2-His protein purified from HEK293T conditioned medium. **(F)** Representative images of HA-hDAP12-mCherry expressed in HeLa cells. Scale bar: 20 µm. **(G)** HA signal intensities at the plasma membrane were quantified and normalized to mCherry fluorescence to assess relative surface expression levels. Control, n=35; sTREM2, n=35. Data are presented as mean ± SEM. Student’s *t*-test.

### sTREM2 reduces phagocytic activity

Since sTREM2 promotes the dissociation of the TREM2–DAP12 complex and reduces the surface expression of DAP12, we hypothesized that sTREM2 acts as a negative regulator of TREM2 signaling. To test this hypothesis, we examined the effect of sTREM2 on the phagocytic activity of BV2 cells. We added fluorescent beads to BV2 cells with or without sTREM2 pretreatment and quantified the internalized beads after 2 hours. Whereas control cells exhibited robust phagocytic uptake, pretreatment with sTREM2 significantly attenuated the internalization of the beads. In contrast, cells cultured at 4°C, a condition that inhibits phagocytosis, showed a marked reduction in bead uptake (Fig. 5A and 5B). These results suggest that sTREM2 suppresses the fundamental phagocytic capacity of BV2 cells. We next investigated whether sTREM2 also inhibits the uptake of fluorescently labeled amyloid β (Aβ). Consistent with our observations using fluorescent beads, sTREM2 pretreatment markedly suppressed the internalization of Aβ. Furthermore, Aβ uptake was completely abolished in cells cultured at 4°C (Fig. 5C and 5D). Collectively, these findings indicate that sTREM2 impairs the microglial clearance of Aβ, likely by interfering with TREM2-mediated phagocytic pathways.

**Figure 5.**
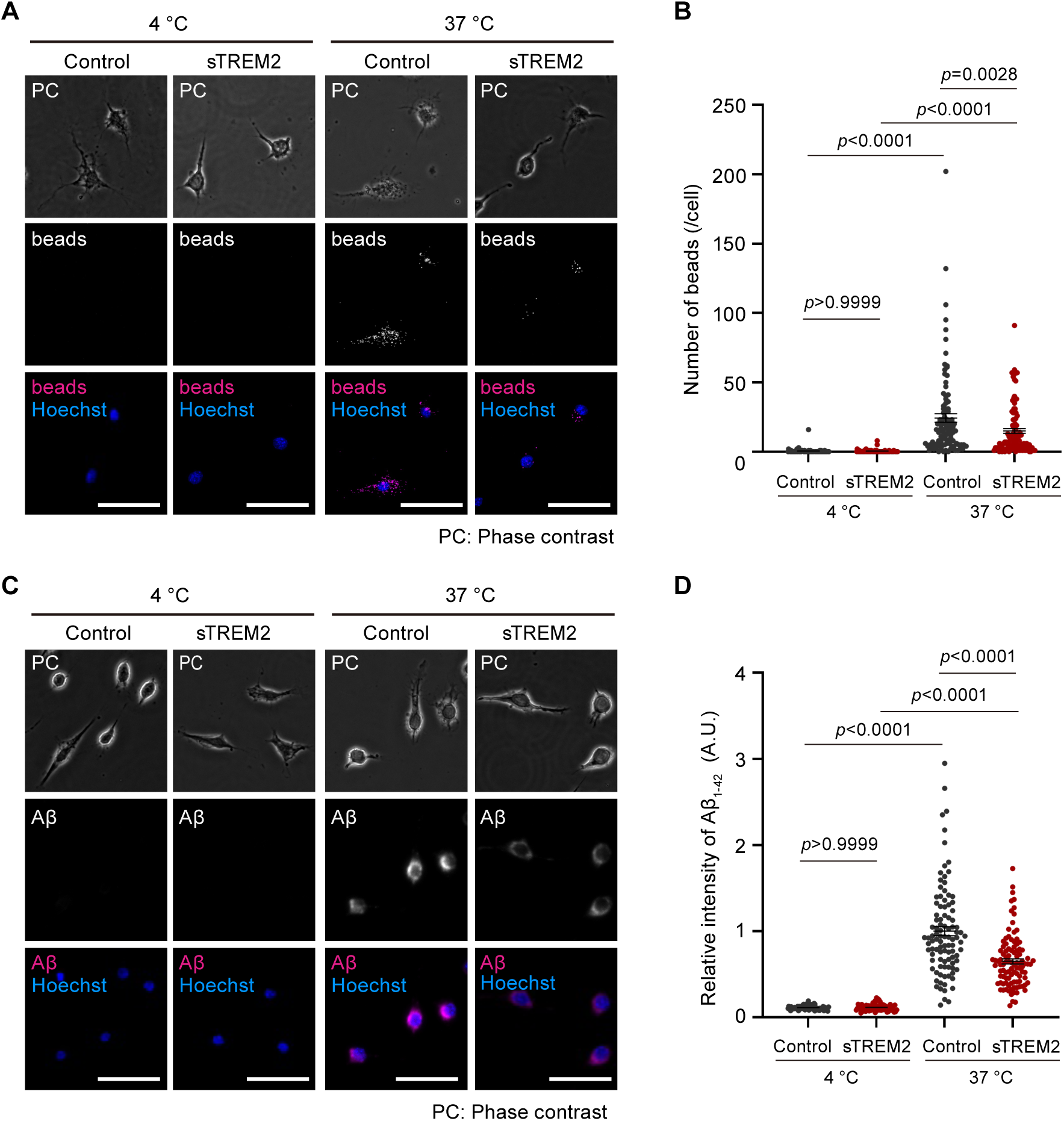
sTREM2 reduces phagocytic activity. **(A)** Representative images of bead uptake in BV2 cells. Cells were pretreated with sTREM2 for 1 h prior to bead incubation. Scale bar, 50 µm. **(B)** Quantification of bead uptake per cell. Control (4 °C), n = 71; sTREM2 (4 °C), n = 81; Control (37 °C), n = 98; sTREM2 (37 °C), n = 106. Data are presented as mean ± SEM. *p* values were analyzed by one-way ANOVA Tukey’s multiple comparisons test. **(C)** Representative images of Aβ uptake in BV2 cells. Cells were pretreated with sTREM2 or control for 1 h prior to incubation with HiLyte Fluor 647–conjugated Aβ₁–₄₂ and then stained with Hoechst. Scale bar, 50 μm. **(D)** Quantification of intracellular Aβ levels. Control (4 °C), n = 85; sTREM2 (4 °C), n = 75; Control (37 °C), n = 100; sTREM2 (37 °C), n = 101. Data are presented as mean ± SEM. *P* values were analyzed by one-way ANOVA Tukey’s multiple comparisons test.

## Discussion

In this study, we demonstrated that sTREM2 interacts with full-length TREM2, promoting the dissociation of the TREM2–DAP12 complex and subsequently reducing the surface expression of DAP12. Furthermore, we found that sTREM2 treatment suppresses the phagocytic activity of the microglial cell line BV2. These findings indicate that sTREM2 modulates microglial function by antagonizing TREM2 signaling, suggesting a novel regulatory mechanism for TREM2-mediated immune responses (Fig. 6A).

**Figure 6.**
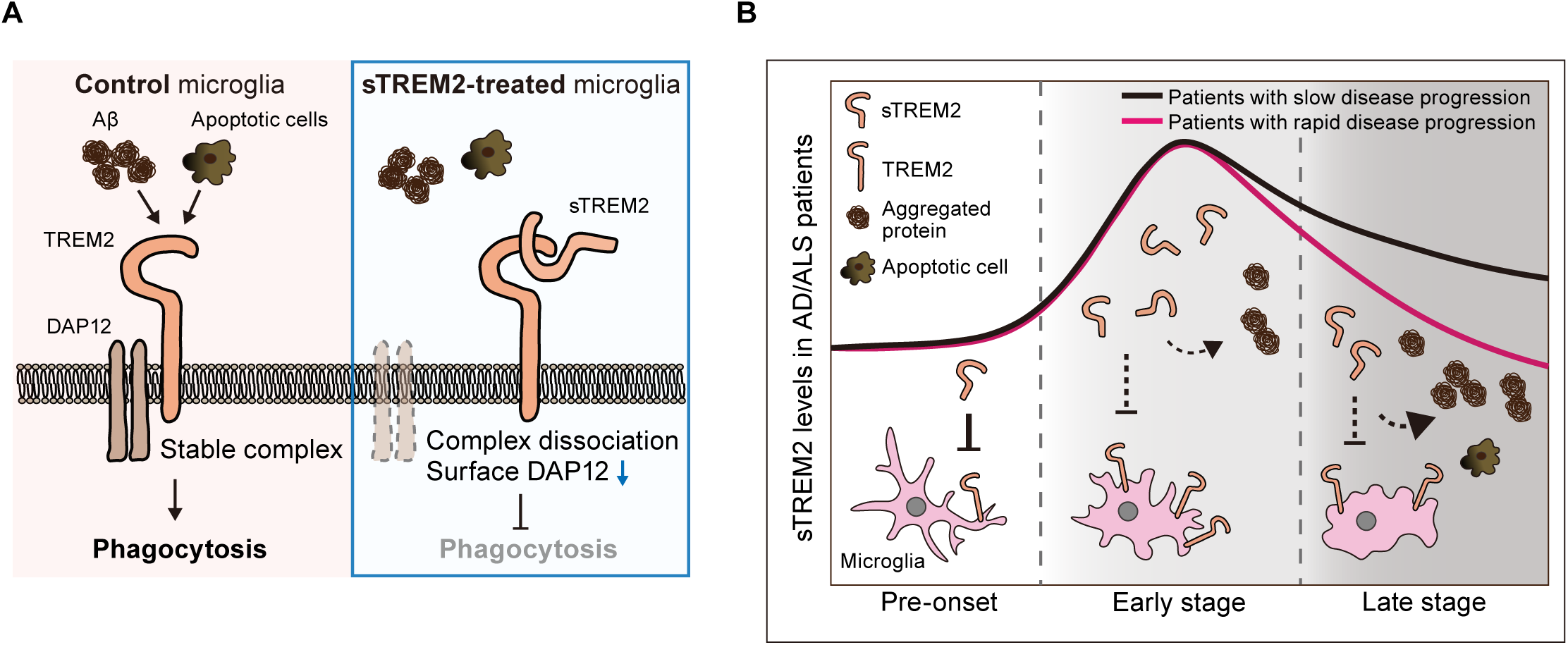
Model for the molecular mechanism of sTREM2 in neuropathological regulation. **(A)** Model of sTREM2-mediated inhibition of TREM2 signaling. sTREM2 antagonizes microglial functions by destabilizing the TREM2–DAP12 complex, leading to an attenuated response to extracellular ligands. **(B)** In neurodegenerative disease, sTREM2 levels in the CSF typically rise during the early stages before gradually declining. A slower rate of sTREM2 reduction is correlated with delayed brain atrophy and reduced protein aggregation. sTREM2 may exert a protective effect in the later stages of the disease, slowing down the progression of symptoms and brain damage.

TREM2 is a critical microglia-enriched receptor that orchestrates microglial responses to neurodegenerative pathologies, including AD and amyotrophic lateral sclerosis (ALS). In AD, TREM2 functions as a sensor for Aβ and lipids, mediating microglial uptake and degradation of Aβ (18, 37). Consistent with these functions, increasing full-length TREM2 on the cell surface by genetic or antibody-mediated blockade of its cleavage reduces amyloid pathology in mouse models of AD (38–40). In ALS models, microglial TREM2 has been shown to contribute to the clearance of pathological protein aggregates, such as TDP-43 and poly-GA (41, 42). These findings suggest that TREM2 plays a protective role in limiting disease progression in neurodegenerative disorders.

However, accumulating evidence indicates that the role of TREM2 is not uniformly protective and can shift toward exacerbating pathology in a context-dependent manner. For instance, studies in tau transgenic mouse models have shown that TREM2 deficiency attenuates neuroinflammation and other pathological features at late disease stages (43). Similarly, TREM2 deficiency in a late-onset AD mouse model accelerates early amyloid plaque seeding but, paradoxically, reduces plaque accumulation at later stages (44). These findings suggest that the impact of TREM2 depends on disease stage. One possible explanation lies in the two-step transition to disease-associated microglia (DAM): the initial activation of microglia proceeds independently of TREM2, whereas the subsequent transition to the fully activated DAM state requires TREM2 signaling(24). Because this TREM2-dependent step coincides with the progression of pathology, sustained TREM2-dependent activation at later stages may promote neuroinflammation and neuronal damage (24, 43, 44). Excessive or inappropriate microglial phagocytosis has also been proposed as a mechanism contributing to neuronal and synaptic loss under neurodegenerative conditions (45).

sTREM2 is generated by proteolytic cleavage of full-length TREM2 and is thought to reflect the microglial response. Consistent with the stage-dependent role of TREM2, cerebrospinal fluid (CSF) sTREM2 levels in AD and ALS patients are elevated at early stages and decline from the middle to late stages (46–48). Higher sustained levels of sTREM2 have been associated with a slower progression of disease pathology and clinical symptoms in AD and ALS patients, suggesting a potential protective role of sTREM2 (48, 49), raising the possibility that the late-stage decline in sTREM2 contributes to disease progression. This appears to conflict with our finding that sTREM2 suppresses phagocytosis in BV2 cells. However, if sustained TREM2-dependent activation becomes detrimental at later stages, sTREM2 may act as a brake that restrains excessive TREM2 signaling and limits neuroinflammation and neuronal damage. In contrast, previous studies have reported that sTREM2 promotes microglial survival and phagocytosis (32, 50). This discrepancy may reflect interactions of sTREM2 with factors other than TREM2. We therefore propose that stage-dependent changes in sTREM2 levels modulate TREM2 function through distinct mechanisms, thereby shaping disease progression (Fig. 6B).

TREM2 variants substantially modulate the risk of neurodegenerative disorders, with specific amino acid substitutions associated with distinct clinical phenotypes. The most extensively characterized variant, R47H, which is located within the extracellular Igl domain, is a strong genetic risk factor for late-onset AD (51, 52). This substitution markedly reduces the binding affinity of TREM2 for Aβ and lipids, thereby impairing microglial plaque clustering and potentially accelerating neurodegeneration (18, 21, 37, 53). Similarly, the R62H variant confers an increased AD risk, albeit with a more modest effect size than R47H, suggesting the functional importance of arginine residues in ligand recognition (51, 54). Moreover, homozygous T66M and Y38C variants are classically associated with Nasu–Hakola disease, a rare disorder characterized by early-onset dementia and systemic bone cysts (55). These mutations commonly lead to protein misfolding and defective trafficking, causing retention of TREM2 within the endoplasmic reticulum and the loss of mature protein at the cell surface (34, 53). Notably, T66M and Y38C variants have also been identified in patients with frontotemporal dementia-like phenotypes without skeletal abnormalities (56). This observation supports a model in which TREM2 dysfunction lies along a biological continuum: partial loss-of-function variants, such as R47H, increase susceptibility to late-onset neurodegeneration, whereas severe structural disruption of the protein, as seen with T66M and Y38C, causes severe, early-onset pathology (26, 57). A detailed understanding of these variant-specific effects will be essential for the development of precision therapeutic strategies to restore TREM2-dependent microglial homeostasis.

A previous study reported that human sTREM2 is monomeric in solution (58). However, our co-immunoprecipitation experiments showed that sTREM2 binds to full-length TREM2 with both mouse and human proteins (Fig. 2A, 2B, 4C and 4D). In addition, the extracellular region of TREM2 multimerizes in molecular dynamics simulations (59), which is consistent with the possibility that sTREM2 binds full-length TREM2. sTREM2 may exist in a variety of states, such as monomers, dimers, and multimers. Also, a monomeric state of purified sTREM2 in solution does not preclude its binding to full-length TREM2, because the high local concentration and clustering of the receptor in the membrane may favor the interaction. Defining the stoichiometry and binding affinity of the sTREM2–TREM2 interaction using purified proteins, for example by size-exclusion chromatography coupled with multi-angle light scattering and surface plasmon resonance, will be important in future studies.

In conclusion, our findings suggest a novel mechanism of TREM2 signaling regulation and provide new insight into how microglial function is controlled in neurodegenerative diseases. Further studies in vivo and in disease models will be important to clarify the role of sTREM2 in disease progression and to evaluate its potential as a therapeutic target.

## Experimental Procedures

### Plasmids

The mouse Trem2 cDNA was amplified by polymerase chain reaction (PCR) from BV2 cDNA library and inserted into the BglII site of pCS4, pCS4-HA, pCS4-His, pCS4-Flag, and pCS4-mCherry plasmids, respectively. The mouse Tyrobp cDNA was also amplified from BV2 cDNA library and inserted into the BglII and XbaI sites of pCS4, pCS4-HA, pCS4-Flag, and pCS4-mCherry plasmids, respectively. The sTrem2 cDNA was amplified from full length Trem2 and subcloned into BglII sites of pCS4-HA and pCS4-His plasmids. The human TREM2 cDNA was amplified by PCR from human fetal brain cDNA library (Dualsystems Biotech, #P12402) and inserted into the BglII site of pCS4, pCS4-His, and pCS4-Flag plasmids, respectively. The human TYROBP cDNA was also amplified from human fetal brain cDNA library and inserted into the BglII and XbaI sites of pCS4-Flag and pCS4-mCherry plasmids, respectively. The human sTREM2 cDNA was amplified from human fetal brain cDNA library and inserted into BglII sites of pCS4-HA and pCS4-His plasmids. To insert the HA tag into the pCS4-Trem2-mCherry and pCS4-Tyrobp-mCherry plasmids, linear plasmids containing the Trem2-mCherry and Tyrobp-mCherry sequences were generated from the respective templates by inverse PCR, followed by ligation of the amplified products. The Trem2 K183A mutant plasmids were generated from the Trem2 plasmids using site-directed mutagenesis and were then circularized using an XthA-ExoT cocktail (60). To construct the pCS4-Trem2 (ΔIgl)-Flag plasmid, the pCS4-Trem2-Flag vector was digested with PstI and then recircularized by self-ligation. The cDNA of Trem2-Igl was amplified from inverse PCR using pCS4-Trem2-His plasmid and ligated using an XthA-ExoT cocktail. After constructing pCS4-Trem2-Igl-His plasmid, the insert was digested using HindIII and XbaI and inserted into pCS4-HA and pCS4-Flag plasmids. The primers and corresponding templates used for PCR are listed in Supplementary Table 1.

### Antibodies

The following primary antibodies were used for immunoblotting: rabbit anti-His antibody (1:1000; MBL, #PM032), sheep anti-TREM2 antibody (1:1000; R&D Systems, #AF1729), rabbit anti-DAP12 antibody (1:1000; Cell Signaling Technology, D7G1X, #12492), mouse anti-Flag antibody (1:1000; Sigma-Aldrich, M2), and rat anti-HA antibody (1:1000; Roche, 3F10). Horseradish peroxidase (HRP)-conjugated secondary antibodies against rabbit, mouse, sheep, and rat IgG (1:20000; SeraCare) were used to detect immunoreactive bands. For immunofluorescence, the primary antibodies used were rat anti-HA (1:1000; Roche, 3F10). Alexa Fluor 488-conjugated anti-rat IgG antibodies (Thermo Fisher Scientific) were used as secondary antibodies.

### Cell culture and transfection

HEK293T and HeLa cells were maintained in Dulbecco’s modified Eagle’s medium (DMEM, high glucose) (FUJIFILM Wako) supplemented with 5% fetal bovine serum (FBS) and 1% penicillin–streptomycin (Thermo Fisher Scientific) at 37 °C in an incubator with 5% CO₂ (61). BV2 cells were maintained in DMEM/F12 (Thermo Fisher Scientific) supplemented with 10% FBS and 1% penicillin–streptomycin under the same conditions (62). For transfection, cells were plated 1 day before transfection at the following densities: HEK293T cells at 2.5 × 10⁶ cells per 150 mm dish (for protein purification) or 8 × 10⁵ cells per 60 mm dish (for immunoprecipitation); HeLa cells at 2 × 10⁴ cells per well in 12-well plates (for immunocytochemistry) or 3 × 10⁵ cells per 35 mm dish (for cell-surface biotinylation assay). Cells were transfected using polyethyleneimine MAX (PEI Max) (Polysciences). Plasmid DNA and PEI Max were mixed in Opti-MEM and added to the cells. After 1 h, the medium was replaced with fresh culture medium. Cells were further cultured for 24 h prior to analysis. For protein purification, cells were subsequently cultured in serum-free medium for an additional 72 h.

### Purification of recombinant protein

HEK293T cells were transfected with an sTREM2-His expression plasmid and cultured in the absence of serum for 72 h. The conditioned medium was collected and centrifuged at 3,000 × g for 5 min to remove cell debris. The resulting supernatant was passed through a 0.22 μm filter. This filtered medium was then loaded onto a column packed with Ni-affinity resin (GE Healthcare, Ni Sepharose 6 Fast Flow), washed sequentially with binding buffer (20 mM Tris-HCl (pH 8.0), 500 mM NaCl, 5 mM imidazole) and wash buffer (20 mM Tris-HCl (pH 8.0), 500 mM NaCl, 60 mM imidazole). The sTREM2 protein was eluted using elution buffer (20 mM Tris-HCl (pH 8.0), 500 mM NaCl, 400 mM imidazole). Finally, the eluate was dialyzed against 1 L of cold phosphate-buffered saline (PBS) at 4°C overnight. The protein concentration was determined by the Bradford assay, and a total of approximately 80 μg of protein was obtained.

### Immunoprecipitation

HEK293T cells were washed with PBS and collected with lysis buffer (50 mM Tris–HCl (pH 7.5), 150 mM NaCl, 0.5% NP-40, 5 mM EDTA). The lysates were centrifuged at 18,000 x g for 10 min at 4 °C, and the supernatants were collected. The supernatants were mixed with either 2 μg of anti-Flag antibody (Sigma-Aldrich, M2) with a 50% slurry of Protein A Sepharose (GE Healthcare, Protein A Sepharose beads 4 Fast Flow) or 0.1 μg of anti-HA antibody with a 50% slurry of Protein G Sepharose beads (GE Healthcare, Protein G Sepharose 4 Fast Flow) with rotation at 4 °C for 3 h. The beads were washed with lysis buffer and subjected to immunoblot analysis.

### Immunoblot analysis

Samples were mixed with 4x sample buffer (50 mM Tris–HCl (pH 6.8), 1.6% SDS, 8% Glycerol, 0.04% bromophenol blue, 20% 2-mercaptoethanol) and boiled at 98 °C for 3 min. Proteins were separated by SDS–polyacrylamide gel electrophoresis and transferred to a polyvinylidene difluoride (PVDF) membrane (Pall). Membranes were blocked with 5% skim milk in Tris-buffered saline with 0.05 % Tween 20 (TBST) for 1 h at room temperature, followed by incubation with primary antibodies overnight at 4 °C. After washing with TBST, membranes were incubated with HRP-conjugated secondary antibodies. Protein signals were detected using a chemiluminescent substrate (Chemi-Lumi One Super, Nacalai Tesque).

### Cell-surface biotinylation assay

BV2 cells were treated with or without sTREM2 (400 ng/mL) for 1 h prior to surface labeling. HeLa cells transfected with the indicated plasmids were not treated with sTREM2. The cells were then washed with PBS and incubated with 0.25 mg/mL EZ-Link Sulfo-NHS-SS-Biotin (Pierce, #A39258) in PBS for 30 min at room temperature. After labeling, cells were washed once with TBS (50 mM Tris–HCl, 150 mM NaCl) to quench unreacted biotinylation reagent and remove excess biotin and twice with PBS and collected with lysis buffer. The lysates were centrifuged at 18,000 x g for 10 min at 4 °C, and the supernatants were collected. The supernatants were mixed with Streptavidin Sepharose (GE Healthcare, Streptavidin Sepharose High Performance) and rotated at 4 °C for 3 h. The beads were washed with lysis buffer and subjected to immunoblot analysis.

### Immunocytochemistry

HeLa cells were transfected with the indicated constructs and incubated for 24 h. Cells were then treated with sTREM2 (400 ng/mL) for 1 h. Cells were fixed with 4% paraformaldehyde (PFA) in PBS for 10 min at 4°C. HeLa cells on coverslips were blocked in blocking solution (5% bovine serum albumin [BSA] in PBS) with or without 0.4% Triton X-100 for 30 min at room temperature, and were then incubated with HA antibody (1:1000) diluted in blocking solution for overnight at 4 °C. After washing with PBS, the cells were incubated with anti-rat Alexa Fluor Plus 488 (1:1000) in blocking solution for 60 min at room temperature. Nuclei were stained with 10 μg/mL Hoechst 33342. The coverslips were then mounted onto slides using FLUOROSHIELD Mounting Medium (ImmunoBioScience). Fluorescent images were obtained using a fluorescence microscope (BZ-X1000, Keyence) with 100× (PlanApo VC 100×/1.4 Oil) objective and a charge-coupled device camera (Keyence).

### Structural analyses

The soluble ectodomains of mouse (UniProt Q99NH8) and human (UniProt Q9NZC2) TREM2 comprised residues 19–157, from the end of the signal peptide (aa 1–18) to His157, immediately before the ADAM10/17 cleavage site. Monomeric structures of the mouse and human sTREM2 were predicted using AlphaFold 3 through the AlphaFold Server (63) with default settings and PDB templates enabled (model seeds 1531826232 for human and 1033229644 for mouse; five models per run). The top-ranked model from each prediction was used for visualization (ranking score, 0.82 and 0.83; pTM, 0.74 and 0.75 for mouse and human, respectively). Igl domains of mouse and human models were superimposed by least-squares fitting (64) of the 112 corresponding Cα atoms using the pair_fit command in PyMOL. Structural similarity was evaluated by the Cα root-mean-square deviation (RMSD) calculated over the same atoms and by TM-score using TM-align (65). TM-score was normalized to the length of the human Igl domain. To assess robustness to model uncertainty, Cα RMSD of the Igl domains was calculated for all 25 pairwise combinations of the five human and mouse models. As a reference for the intrinsic variability of the predictions, Cα RMSDs were also calculated for all pairwise combinations among the five human models and among the five mouse models. Structures were rendered using open-source PyMOL 3.1 (Schrödinger, LLC) with ray tracing on a white background. The mature sequences were aligned without gaps. Identical, strongly similar and weakly similar positions were defined by the Clustal conservation groups. Secondary structure shown in Fig. 4A was assigned with the PyMOL dss command for the top-ranked human model. Computational analyses were performed in Python 3.11 using Biopython 1.88 (66), gemmi 0.7.5, NumPy 2.4, SciPy 1.17, tmtools 0.3.0, and Matplotlib 3.11.**s**

### Phagocytosis assay

BV2 cells were plated at 2.5 × 10⁴ cells per well in 24-well plates and cultured for 1 day. For beads engulfment assay, BV2 cells were pretreated with sTREM2 (400 ng/mL) for 1h, washed with fresh medium, and then incubated with 1 μL/mL Fluoresbrite Polychromatic Red Microspheres (0.5 um; Polysciences, Inc., #19507-5) for 2 h at 37 °C. As a negative control, cells were incubated at 4 °C to inhibit phagocytosis. After incubation, cells were washed with PBS with gentle agitation to remove non-internalized beads and surface-bound beads, and fixed with 4% paraformaldehyde in PBS for 10 min at 4 °C. Cells were then washed with PBS and blocked with 5% BSA in PBS. Nuclei were stained with 10 μg/mL Hoechst 33342. The coverslips were then mounted onto slides using FLUOROSHIELD Mounting Medium (ImmunoBioScience). Fluorescence and phase contrast images were acquired using a microscope (BZ-X1000, Keyence) with 40x (PlanFluor NA 40x/0.60) objective and a charge-coupled device camera (Keyence). For Aβ engulfment assay, Phagocytosis of aggregated HiLyte Fluor 647–conjugated Aβ₁-₄₂ (Anaspec, #AS-64161) was analyzed as previously described with minor modifications (34, 67, 68). Aβ₁-₄₂ was prepared as follows. 0.1 mg of peptide was dissolved in 40 μL of hexafluoroisopropanol (HFIP) and incubated for 30–60 min at room temperature. After evaporation of HFIP, the peptide was resuspended in dimethyl sulfoxide (DMSO). The prepared Aβ₁-₄₂ was then aggregated overnight at 37 °C with agitation. BV2 cells were pretreated with sTREM2 (400 ng/mL) for 1h, washed with fresh medium, and then incubated with 500 nM aggregated Aβ₁-₄₂ for 3 h at 37 °C. As a negative control, cells were incubated at 4 °C to inhibit phagocytosis. Extracellular Aβ₁-₄₂ was quenched with 0.2% trypan blue in PBS for 1 min. Cells were washed with PBS and fixed with 4% paraformaldehyde in PBS for 10 min at 4 °C. BV2 cells were then blocked with 5% BSA in PBS. Nuclei were stained with 10 μg/mL Hoechst, and coverslips were mounted. Fluorescence and phase contrast images were acquired using a microscope (BZ-X1000, Keyence) with 40x (PlanFluor NA 40x/0.60) objective and a charge-coupled device camera (Keyence).

### Statistical Analyses

The statistical analyses were calculated by using GraphPad Prism v.10.4.1 (GraphPad Software). All image processing and quantifications were conducted by FIJI Image J v1.54p. For comparisons between two groups, an unpaired two-tailed Student’s t-test was used. For comparisons among more than two groups, one-way ANOVA followed by Tukey’s multiple comparisons test was used. All data were reproduced in at least three independent experiments.

## Acknowledgments

We thank the lab members for helpful discussions and technical support. This work was supported by AMED-PRIME [ADI07313 (FT)], Gout and uric acid Foundation (FT), Sumitomo Foundation (FT), Ono Pharmaceutical Foundation for Oncology, Immunology, and Neurology (FT), and Center for Quantum and Information Life Sciences, University of Tsukuba (FT).

